# Development of fluorescent peptide G Protein Coupled Receptor activation biosensors for NanoBRET characterisation of intracellular allosteric modulators

**DOI:** 10.1101/2022.05.30.494024

**Authors:** James P Farmer, Shailesh N Mistry, Charles A Laughton, Nicholas D Holliday

## Abstract

G protein coupled receptors (GPCRs) are widely therapeutically targeted, and recent advances in allosteric modulator development at this class of receptors offer further potential for exploitation. In particular GPCR intracellular allosteric modulators (IAM) represent a class of ligands that bind to the receptor-effector interface (e.g. G protein) and so inhibit agonist responses non-competitively. This potentially offers a tailored mode of action and greater selectivity between conserved receptor subtypes compared to classical orthosteric ligands. However, while specific examples of the IAM class of ligands are well described (particularly for chemokine receptors), a more general methodology for assessing compound interactions at the GPCR IAM site is lacking. Here fluorescent labelled peptides based on the Gα peptide C terminus are developed as novel binding and activation biosensors for the GPCR IAM binding site. In TR-FRET binding studies, unlabelled peptides derived from the GαS subunit C-terminus were first characterised for their ability to positively modulate agonist affinity at the β_2_-adrenoceptor. On this basis, a tetramethylrhodamine (TMR) labelled tracer was synthesized based on the 19 amino acid C terminal GαS peptide (TMR-GαS19cha18, where cha=cyclohexylalanine). Using NanoBRET technology to detect binding, TMR-GαS19cha18 was recruited to Gs coupled β_2_-adrenoceptor and EP2 receptors in an agonist dependent manner (correlated with ligand efficacy), but not to the Gi coupled CXCR2 receptor. Moreover, NanoBRET competition binding assays using TMR-GαS19cha18 enabled direct assessment of the affinity of unlabelled ligands for β_2_-adrenoceptor IAM site. Thus the NanoBRET platform using fluorescent-labelled G protein peptide mimetics offers novel potential for medium-throughput affinity screens to identify new IAMs, applicable across GPCRs coupled to a G protein class. Using the same platform, Gs peptide biosensors also represent useful tools to probe orthosteric agonist efficacy and the dynamics of receptor activation.

## Introduction

G protein coupled receptors (GPCRs) are the largest family of membrane bound receptors and, with over 27% of the global market share for therapeutic drugs, continue to be the most clinically targeted amongst receptor superfamilies^1^. Within the GPCR superfamily, the class A rhodopsin like receptors are most numerous (approximately 350 members), and are grouped by conservation of key amino acid motifs supporting the seven transmembrane helical bundle structure and conformational changes on activation^1^. Classical GPCR signalling involves the formation of an agonist-receptor-effector complex (the allosteric ternary complex^2^) to activate heterotrimeric G proteins (or other signalling proteins, such as b-arrestins). Class A GPCRs primarily signal through Gαs, Gαi/o, Gαq/11 or Gα12/13 containing G proteins, each with distinct functional outcomes including cAMP and calcium mobilisation, modulation of ion channel activity and protein kinase cascades and gene expression ^3,4 5,6^. A key component of the interaction between the activated GPCR and the Gα subunit is determined by the Gα C terminus (the α5 helix), which engages the GPCR intracellular domain in a cleft between helices TM3 and TM5^2,7–9^. The sequence of the Gα α5 helix is an important driver determining the selectivity of GPCR interaction with different Gα classes; its selective interaction with the active agonist-bound receptor conformation provides the basis for the ternary complex and allosteric stabilization of agonist high affinity binding to the receptor in the presence of the G protein6.

The nature of the GPCR-G protein interaction pocket provides opportunity for small molecule intervention to modulate receptor function, distinctive from classical targeting of the orthosteric binding site. Indeed, high affinity intracellular allosteric modulators (IAMs) have been reported that act as non-competitive antagonists to prevent signalling, for a range of Gi and Gs coupled receptors including the chemokine receptors CCR2, CXCR2, CCR5 and CCR9; the β_2_-adrenoceptor (β_2_-AR); and most recently the prostanoid receptor EP2^8,10–14^. In line with other approaches to develop allosteric GPCR ligands^8,15^, the rationale for IAM development includes the ability to develop selective ligands against receptor subtypes with conserved orthosteric binding sites, or ones which are multifaceted and complex for small molecule design (e.g. chemokines). Intracellular allosteric modulation may also give rise to useful pharmacological properties – for example, an IAM series at the EP2 prostanoid receptor displayed use dependence, arising from their higher affinity for the activated agonist-GPCR conformation, while maintaining blockade of receptor-effector coupling^14^. However, to date, a general target methodology is lacking for identifying novel IAMs by their affinity for the target GPCR-G protein interface.

Early influential studies to explore the receptor-G protein coupling mechanism demonstrated the particular role of the Gα C terminal α5 helix through the use of peptide mimetics or expressed mini-genes, showing their ability to compete with native G protein binding and inhibit GPCR signaling depending on their sequence^16^,^17–20^. However further exploitation of these peptides as tools was initially limited by relatively modest affinity, ways to assess direct engagement of the peptides with the receptor, and issues such a lack of cell permeability for functional studies. Recently Mannes et al (2021)^21^ identified a number of modifications to GαS C terminal peptides capable of improving affinity for the β_2_-adrenoceptor, and also demonstrating their use dependent properties in which the high agonist affinity active conformation of the GPCR was selectively stabilised. Given that activity was preserved at another Gs coupled GPCR (the D1 dopamine receptor)^21^, these peptides therefore offer starting points for broad selectivity tool development for assessing GsPCR IAM binding sites.

Increasingly, the use of fluorescent ligands and resonance energy transfer technologies (such as TR-FRET and NanoBRET) provides an attractive alternative to radioligand and other approaches to assess binding in both GPCR mechanism of action studies and compound screening ^22–26^. The selectivity of the resonance energy transfer signal (constrained to a distance of less than 10 nm between the donor luciferase / fluorophore and acceptor fluorescent tracer) enables these assays to be performed in a homogeneous format (without separation of the free fluorescent ligand) and accurate determination of ligand binding even at high tracer concentrations. In this study we demonstrate a novel generally applicable NanoBRET approach to monitor the binding of ligands at GPCR IAM sites, to quantify interactions between inactive / activated GPCRs and a fluorescent derivative of the 19 amino acid GαS C terminal peptide GαS19cha18 (where cha=cyclohexylalanine)^21^. Supported by assessments at both the β_2_-adrenoceptor and EP2 prostanoid receptor, we show how this provides a sensitive real time biosensor for receptor activation by agonists of differing efficacy, and generally applicable binding assay for determination of the ligand affinity at the GPCR-G protein IAM binding site.

## Materials and Methods

### Materials

Gα C terminal peptides were purchased from GenScript Biotech (New Jersey, USA) (4mg, >95% purity) and stored at -20°C at a 10mM stock concentration in DMSO prior to use (sequences given Table 1). General molecular biology enzymes were obtained from Fermentas (ThermoFisher Scientific, Loughborough, UK), and other molecular biology consumables were from Sigma-Aldrich (Poole, UK) or ThermoFisher (Loughborough, UK) unless otherwise stated. All assay plates used were OptiPlate-384 white well microplates (product number: 6007290, PerkinElmer LAS Ltd, UK) unless otherwise stated. BODIPY-FL-PEG8-(S)-Propranolol was purchased from Hello Bio Ltd (CA200693, Bristol, UK) and all other compounds were purchased from ThermoFisher (Loughborough, UK) and stored as 10mM stocks in DMSO at -20°C unless otherwise stated. The nanoluciferase substrate furimazine was purchased from Promega Biotech (Madison, USA).

**Table 1:** Amino acid sequences of GαC terminal peptides used in this study

| Gα C-terminal peptide | Amino acid sequence (N to C terminus left to right) |
| --- | --- |
| GαS11 | QRMHLR QYELL |
| GαS24 | NIRRVFNDCRDII QRMHLRQYELL |
| GαS19cha18 | FNDCRDII QRMHLRQYE{CHA}L |
| TMR-GαS19cha18 | **TMR**-FNDCRDII QRMHLRQYE{CHA}L |
| Gαi24 | NVQFVFDAVTDVI IKNNLKDCGLF |

{CHA} indicates cyclohexylalanine, **TMR** indicates N terminal incorporation of Tetramethylrhodamine

### Cell culture

HEK 293 cell lines were transfected with cDNAs encoding (i) N-terminal SNAP-tagged human β_2_-adrenoceptor (Hek-ssβ_2_-AR, Genbank: NM_000024, as described in Valentin-Hansen et al (2012))^27^ or (ii) SNAP-tagged human β_2_-adrenoceptor or human EP2 receptors fused at the C terminus to a thermostable Nanoluciferase (Hek-ssβ_2_-AR-tsNluc) using Lipofectamine 3000 reagent (Invitrogen, Paisley, UK). Original SNAP tag and Nanoluciferase sequences were from NEB (Hitchen UK) and Promega (Southampton UK) respectively. The tsNluc contains structural stabilising Nluc substitutions as described in Hoare et al^28,29^. Stable mixed population cell lines were established through G418 resistance (encoded by the plasmid vector (pcDNA3.1 neo^+^, Invitrogen, Paisley UK). Cells were maintained in Dulbecco’s modified Eagle’s medium (Sigma-Aldrich, Poole, UK) supplemented with glucose (4.5g/L), 0.2mg/ml G418, L-Glutamine (4.5g/L), and with 10% fetal calf serum (Life Technologies, Paisley, U.K.)

### Terbium Labelling of SNAP-tagged human β_2_-adrenoceptors, and membrane preparations

For TR-FRET assays requiring terbium (Tb) labelling, cell culture medium was removed from T-175cm^2^ flasks containing confluent adherent Hek-ssβ_2_-AR and replaced with 10ml Tag-lite medium (LABMED, Cisbio Bioassays) containing 100nM SNAP-Lumi4-Tb (Cisbio Bioassays, Bagnols-sur-Ce’ze,France). Cells were then incubated in labelling medium for one hour at 37°C under 5% CO_2_. For membrane preparations, Tb-labelled Hek-ssβ_2_-AR cells and unlabelled Hek-ssβ_2_-AR-tsNluc,HEK-ssEP2-tsNluc, or HEK-ssCXCR2-tsNluc cells were washed twice with phosphate-buffered saline (PBS, Sigma-Aldrich, Pool, UK) to remove excess labelling or growth medium before being removed by scraping into 10ml PBS. Detached cells were then collected and pelleted by centrifugation (10 minutes, 2000rpm) and pellets were frozen at -80°C, until required. For membrane homogenization (all steps at 4°C), 20ml wash buffer (10mM HEPES, 10mM EDTA, pH: 7.4) was added to the pellet before disruption (8 bursts) with an Ultra-Turrax homogenizer (Ika-Werk GmbH & Co. KG, Staufen, Germany), and subsequently centrifugation at 48 000*g* at 4 °C. Supernatant was discarded and the pellet was resuspended in 20ml wash buffer and centrifuged again as above. The final pellet was suspended in cold 10 mM HEPES with 0.1 mM EDTA (pH 7.4). Protein concentration was determined using the bicinchoninic acid assay kit (Sigma-Aldrich, Pool, UK) using bovine serum albumin as standard, and aliquots were maintained at -80 °C until required.

### TR-FRET BODIPY-FL-PEG8-(S)-Propranolol binding assays to determine association kinetics and equilibrium ligand binding

TR-FRET binding assays were performed using low sodium assay binding buffer (25mM HEPES, 1% DMSO, 0.1mg/ml Saponin, 0.02% w/v Pluronic acid F127, 1mM MgCl2 and 0.2% BSA (pH 7.4), at 37°C in 384 well Optiplates. To determine fluorescent ligand association kinetics, incubations were performed BODIPY-FL-PEG8-(S)-Propranolol (Fl-propranolol) at varied final assay concentrations (1.56 - 100nM) in the absence and presence of 10 μM ICI118551 to determine non-specific binding. Binding was initiated by the addition of Hek-ssβ_2_-AR cell membranes (1μg/well) in assay buffer to a final assay volume of 30μl, performed by online injection on a BMG PHERAstar FSX platereader (BMG Labtech, Offenburg, Germany). The TR-FRET ratio was then recorded at 10 s intervals, over a 30 min period, on the PHERAstar using 337 nm excitation of the Terbium donor and monitoring donor emission at 490 nm, and acceptor Fl-propranolol emission at 520 nm.

For TR-FRET competition binding studies in the same system, assays were performed using the same buffer and temperature conditions above, using 20 nM Fl-propranolol tracer in the absence and presence of 14 competing concentrations of unlabelled test orthosteric ligands (salbutamol, isoproterenol, formoterol and ICI118551), with or without 10μM Gα C terminal peptide (GαS11, GαS24, GαS19cha18 or Gαi24; Table 1). Binding was initiated by online addition of the Hek-ssβ_2_-AR cell membranes (1 μg / well) to a final assay volume of 40 μl. Endpoint HTRF readings were taken at 30 – 120 min timepoints using the PHERAstar HTRF settings to monitor progress to equilibrium. For these studies, total binding was determined by using assay buffer in the place of competing ligand and NSB was determined with 10μM ICI118551.

### Bioluminescence resonance energy transfer (NanoBRET) assays to monitor fluorescent G protein peptide recruitment

NanoBRET assays used either low sodium assay binding buffer as described for TR-FRET binding measurements, or an extracellular Hank’s balanced saline solution (136 mM NaCl, 5.1 mM KCl, 0.44 mM KH_2_PO_4_, 4.17 mM NaHCO_3_, 0.34 mM Na_2_HPO_4_, 1% DMSO, 0.1mg/ml Saponin, 0.02% Pluronic acid F_127_, 0.2% BSA and 20mM HEPES, pH 7.4), as indicated in the text. TMR-GαS19cha18 binding was was first characterised by addition of varying fluorescent probe concentrations (8 – 1000 nM) to 1μg/well Hek-ssβ_2_-AR-tsNluc cell membranes, in which donor luciferase luminescence was stimulated with the addition of furimazine (1/960 dilution from Promega manufacturer’s stock) in the additional absence or presence of the orthosteric agonist 10μM isoproterenol (final assay volume, 40μl). Fluorescent peptide NSB was defined by the inclusion of 10μM unlabelled GαS19cha18 peptide. Endpoint reads at 30- and 60-min incubation at 37°C were taken on the PHERAstar as the BRET ratio between donor luminescence (450 nm emission) and acceptor TMR-GαS19cha18 fluorescence (550 nm) to determine TMR-GαS19cha18

For quantitative analysis of TMR-GαS19cha18 recruitment by orthosteric agonists, 500nM TMR-GαS19cha18 was incubated with 1μg/well Hek-ssβ_2_-AR-tsNluc cell membranes, 1/960 dilution furimazine, and the following ligands at the indicated final concentrations: salbutamol, salmeterol, isoproterenol, formoterol, or ICI118551. To initiate the recruitment, membranes were separately preincubated (5 min) with furimazine to establish luminescence output, prior to their online injection using the PHERAstar to assay buffer containing the probe peptide and stimulating ligands.

NanoBRET was monitored for 30 min every 1.16 minutes on the PHERAstar, using the BRET ratiometric measurements described above. In experiments to understand TMR-GαS19cha18 selectivity, agonist binding assays were repeated using the same protocol employing Hek-EP_2_-tsNluc or Hek-ssCXCR2-tsNluc cell membranes, stimulated with PGE2^9^ or CXCL8 (Stratech Scientific, Cambridge, UK), or vehicle. In these experiments the extent of recruitment was assessed 30 min after membrane addition at 37°C.

To determine unlabelled ligand affinities using competition binding, assays employed 500nM or 125nM TMR-GαS19cha18, a range of competing concentrations of unlabelled peptides (e.g., GαS19cha18, GαS24, GαS11 or Gαi24), the inclusion of 10μM isoproterenol and 1μg/well Hek-ssβ_2_-AR-tsNluc cell membranes pre-incubated with 1/960 dilution of furimazine as indicated above (final volume, 50μl). Incubations were performed at 37°C and BRET measurements were taken every 30 minutes over a 2-hour interval, using PHERAstar (550 nm / 450 nm ratio).

### Data analysis

TR-FRET and NanoBRET data were performed in either triplicate or duplicate unless otherwise indicated and were routinely expressed as the respective acceptor / donor emission ratios (520 nm / 490 nm for TR-FRET; 550 nm / 450 nm for NanoBRET) x 1000. In competition binding studies, individual experiment data were normalised to total specific binding in the absence of competing ligands (100 %), while in agonist-stimulated recruitment measurements data were normalised to a maximal concentration of stimulating reference agonist.

For TR-FRET Fl-propranolol association kinetic data, specific binding traces for Fl-propranolol (defined as total binding – NSB) were fitted to a one site association model. Global fitting of this model across multiple fluorescent ligand concentrations from the same experiment enabled estimation of FL-propranolol association (k_on_) and dissociation rate constants (k_off_), together with the kinetically derived K_D_ (=k_off_/k_on_) using the equations:

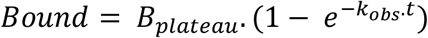

Where, the B_plateau_ is the equilibrium level of tracer binding, and the observed association rate constant k_obs_ is related to the binding rate constants for FL-propranolol in a single site model by:

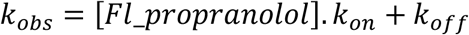

Endpoint saturation analysis also enabled calculation of the equilibrium dissociation constant (K_D_) for fluorescent tracer, as well as total binding density as B_max_ in TR-FRET and BRET experiments, based on:

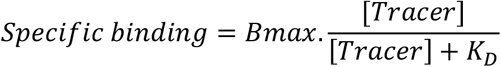

Competiton binding studies were fitted to determine competing ligand IC_50_ concentrations, using a four-parameter fit including the Hill slope (n)

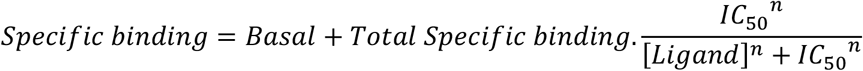

Where appropriate, the Cheng-Prusoff equation was applied to convert IC_50_ estimates to the competing ligand dissociation constant as K_i_

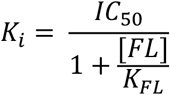

K_FL_ and [FL] represent the fluorescent probe dissociation constant and concentration respectively.

For endpoint agonist stimulation of TMR-GαS19cha18 recruitment measured by NanoBRET, concentration response curve analysis was performed to obtain estimates of ligand potency (EC_50_) and maximal response R_max_:

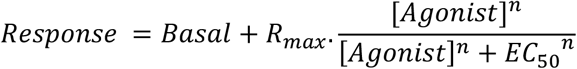

Alternatively kinetic Gs recruitment data were fitted to a rise-to-steady state model, as described by Hoare et al (2020)^27^:

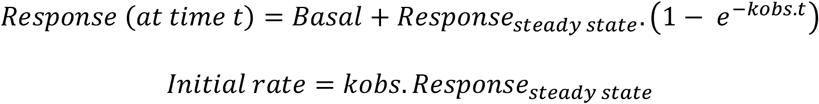

In this analysis, concentration response data were analysed by defining the initial rate at each ligand concentration as the response.

All data analysis was performed using Prism 9.0 (GraphPad Software, San Diego). Parameter estimates were expressed as pX (-log X) where appropriate (e.g. pEC_50_) and data from individual experiments were pooled as mean ± s.e.m. Statistical significance between two data groups was assessed by Student’s unpaired or paired t-test as indicated in the text, with a level of significance defined as p < 0.05.

## Results

### Determination of the effects of GαS C terminal peptides on β_2_-adrenoceptor ligand binding using the BODIPY-FL-PEG8-(S)-Propranolol TR-FRET binding assay

In order to probe the potential allosteric modulatory effects of different Gα C terminal peptides on β_2_-adrenoceptor (β_2_- AR), we first established a membrane TR-FRET binding assay using terbium labelled HEK ssβ_2_-AR membranes and the fluorescent antagonist tracer Fl-propranolol^24,30,31^. The binding parameters of Fl-propranolol were determined through analysis of the association kinetics for the fluorescent tracer (Supplementary Figure 1), confirming single site behaviour and FL-propranolol estimates for the association rate constant k_on_ (1.30±0.18 ×10^7^ M^-1^min^-1^), dissociation rate constant k_off_ (0.18±0.02 min^-1^) and a kinetically derived K_D_ 16.1±3.1 nM (n=5). These estimates are consistent with previously reported data for this ligand at the β_2_-AR^32^.

FL-propranolol competition analysis was then performed to determine the affinities of three representative agonist ligands (isoproterenol, formoterol, salbutamol) and the unlabelled antagonist ICI118551 in a low sodium assay buffer as described in the methods (Figure 1; Table 2) – and to examine the potential allosteric effect of different GαS derived peptides.

**Figure 1:**
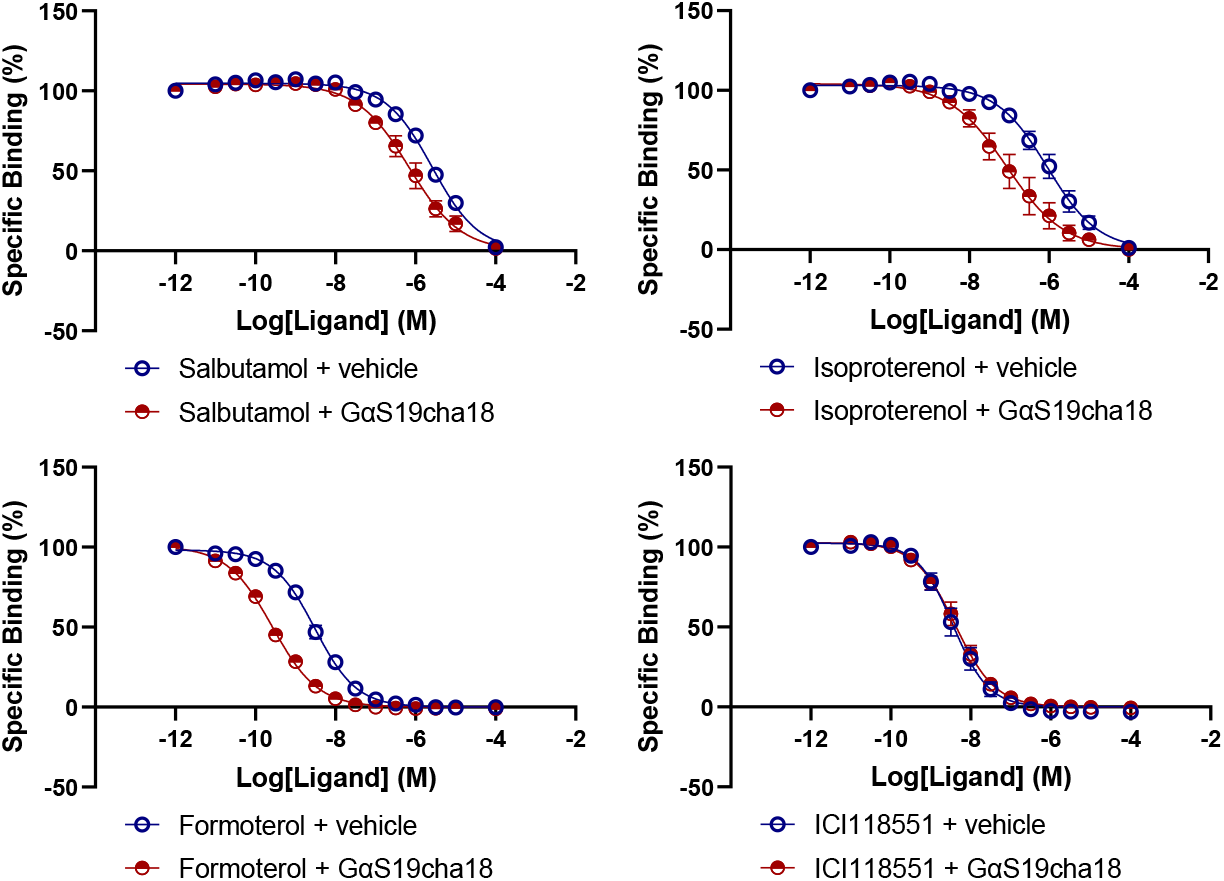
TR-FRET FL-propranolol binding studies in ssβ_2_-AR membranes demonstrate the selective effect of GαS19cha18 (10 μM) on agonist affinities. The specific binding data shown were normalized and pooled from five independent experiments, with non-specific binding was determined by addition of 10μM ICI118551. All assays were performed using 20 nM Fl-propranolol tracer and low sodium buffer at 37°C for 2 hours, with comparator peptide data (GαS24, GαS11) shown in the Supplementary Figures 2 and 3.

**Table 2:**
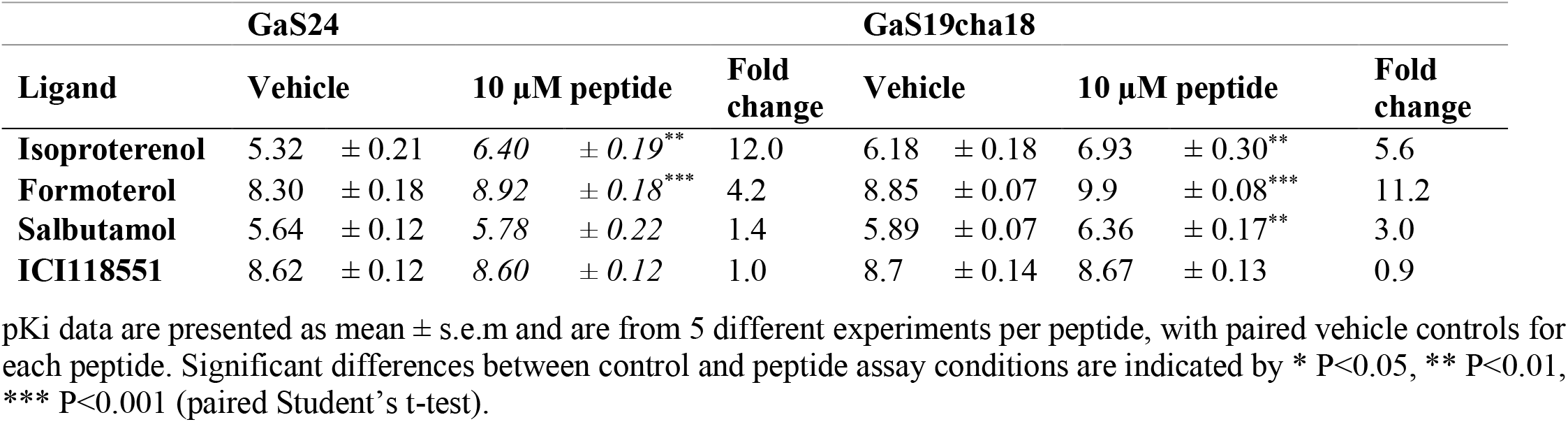
Binding affinities of β2-adrenoceptor ligands in the absence and presence of 10μM Gαs C terminal peptides

Specific binding was monitored at a number of timepoints (data not shown) to ensure the 2 h endpoint shown represented equilibrium conditions. In the absence of GαS peptides, the measured affinity of these example ligands was as expected from literature data^30,32^. Notably 10μM GαS24 and 10μM GαS19cha18 each promoted a significant increase in measured affinity for the orthosteric agonists (Figure 1; Table 2; Supplementary Figure 2). The extent of the shift in affinity was highest for the high efficacy agonists isoproterenol and formoterol and reduced for the lower efficacy agonist salbutamol. In contrast GαS24 and GαS19cha18 had no effect on the affinity of the antagonist ICI118551. We also observed that a shorter 11 amino acid C terminal GαS11 peptide had no effect on orthosteric agonist binding in the same assay, at up to 10μM (Supplementary Figure 3; Supplementary Table 1). Taken together these data supported the binding of GαS24 and GαS19cha18 to the β_2_-AR, and their allosteric stabilisation of the active conformation selectively promoting high agonist affinity, as previously observed in radioligand binding assays by Mannes et al (2021) 23.

### Establishing a NanoBRET assay to directly monitor TMR-GαS19cha18 recruitment to the β_2_-Adrenoceptor

The positive enhancement of agonist binding affinity in the β_2_-AR was greatest for GαS19cha18, supporting previous studies indicating the β_2_-AR affinity and allosteric modulation by this substituted GαS peptide^21^. However, this analysis of the effects of the Gα C terminal peptides relied on their indirect allosteric modulation properties, rather than direct demonstration of binding and peptide affinity for the β_2_-AR-G protein interaction site. Given the knowledge that the peptide C terminus was likely to make close contact with the α5-helix^7^, we therefore sought to generate a fluorescent probe retaining β_2_-AR affinity through N terminal modification of the sequence with the BRET compatible fluorophore tetramethylrhodamine (generating TMR-GαS19cha18). The ssβ_2_-AR was fused at the C terminus with a thermostable (ts) Nanoluciferase (ssβ_2_-AR-tsNluc) thereby providing a source of intracellularly located donor luminescence and providing opportunity to detect TMR-GαS19cha18 to the expressed ssβ_2_-AR-tsNluc in membranes by NanoBRET (Figure 2A).

**Figure 2:**
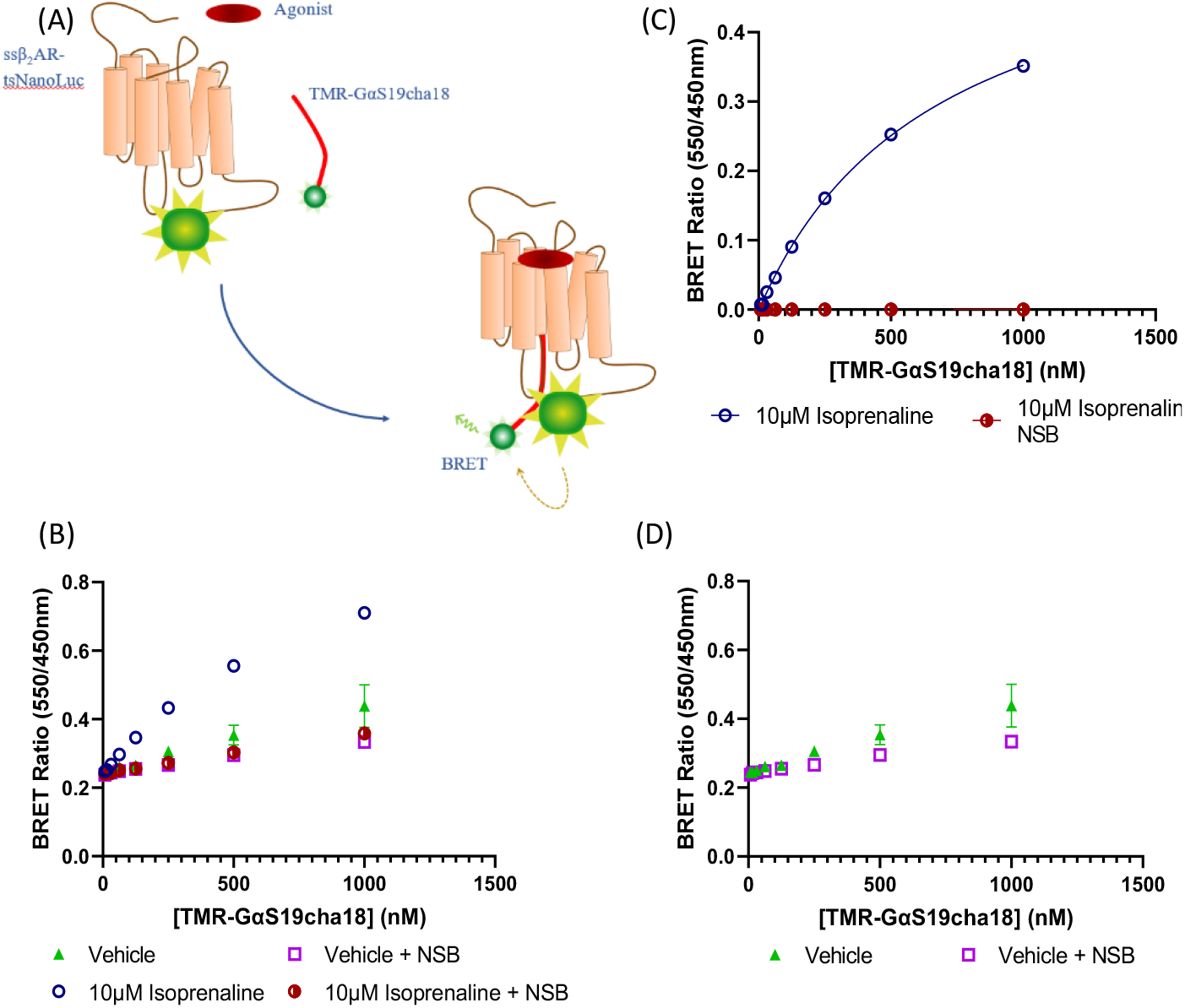
Agonist dependent binding of TMR-GαS19cha18 to the β_2_-adrenoceptor determined by NanoBRET. A) Diagram of TMR-GαS19cha18 interaction with ssβ_2_-AR-tsNluc, generating NanoBRET signal, in the presence of an agonist. B) Saturation binding of TMR-GαS19cha18, demonstrating the increased specific binding observed in the presence of 10μM isoproterenol. C) Specific binding data in the presence of isoproterenol fitted with a one-site specific binding model to determine TMR-GαS19cha18 affinity (K_D_). D) Saturation binding measurements for TMR-GαS19cha18 to ssβ_2_-AR-tsNluc membranes in an agonist free environment and low sodium buffer. In B and D, total and non-specific binding (NSB) were defined by the absence and presence of 10μM unlabelled GαS19cha18. Data shown are single examples from five independent experiments.

However, initial saturation studies performed using TMR-GαS19cha18 and otherwise unstimulated ssβ_2_-AR-tsNluc membranes (in low sodium buffer used for previous TR-FRET measurements) failed to detect significant specific binding using up to 1 μM labelled peptide (Figure 2D). Instead, TMR-GαS19cha18 recruitment was only observed in the presence of 10 μM isoproterenol, in which a substantive specific BRET measurement was observed that was effectively competed by unlabelled GαS19cha18 peptide (Figure 2B, Figure 2C). Under these agonist-stimulated conditions and low sodium environment, the TMR-GαS19cha18 K_D_ for the β_2_-AR was 588 ± 19 nM (n=5). The use of an extracellular HBSS based buffer (with higher sodium concentration) did not significantly affect TMR-GαS19cha18 affinity (Supplementary Figure 4). These data demonstrated that TMR-GαS19cha18 was a suitable probe for the β_2_-AR intracellular G protein binding site, whose binding could be detected by NanoBRET, and appeared dependent on the active receptor conformation promoted by orthosteric β_2_-AR agonists.

### β_2_-Adrenoceptor ligand pharmacology revealed by TMR-GαS19cha18 NanoBRET recruitment assays

Given the agonist dependence of TMR-GαS19cha18 recruitment, we explored the ability of this NanoBRET assay to function as a β_2_-AR activation sensor for ligands of known differences in efficacy. Using 500 nM TMR-GαS19cha18 tracer, kinetic and endpoint NanoBRET measurements were performed in ssβ_2_-AR-tsNluc membranes in response to agonists and antagonist, also comparing the low sodium binding buffer initially used with an HBSS based buffer with higher “extracellular” sodium concentrations. Endpoint concentration response data (Figure 3; Table 3) clearly ranked the agonists isoproterenol, formoterol, salbutamol and salmeterol in the expected order of potency and maximal response^30^, with salbutamol and salmeterol both identified as partial agonists relative to isoproterenol. The effect of HBSS buffer environment, containing higher sodium concentration, was a reduction in agonist potency (Figure 3B, Table 3). A further advantage of the NanoBRET methodology was the homogeneous assay format and the ability to collect the kinetics of TMR-GαS19cha18 recruitment for the different agonists over time (Figure 4). Fitting the rise to steady-state observed in the data enabled calculation of the initial rate of fluorescent G peptide probe recruitment at each agonist concentration^33^, and to construct concentration-initial response rate relationships for the agonists as shown in Figure 4D, Table 4). These data provided equivalent agonist potency and maximal response measurements to the endpoint concentration-response measurements performed under the same buffer conditions.

**Table 3:** Agonist potencies and maximal responses derived from TMR-GαS19cha18 binding in Low sodium buffer or extracellular HBSS media.

| Ligands | Low Sodium |  |  |  | HBSS |  |  |  |
| --- | --- | --- | --- | --- | --- | --- | --- | --- |
|  | pEC<sub>50</sub> | ±s.e.m (M) | R<sub>max</sub> | ±s.e.m (%) | pEC<sub>50</sub> | ±s.e.m (M) | R<sub>max</sub> | ± s.e.m (%) |
| Salbutamol | 6.84 | ±0.16 | 49.2 | ±2.7 | 5.91 | ±0.22** | 26.6 | ±3.4*** |
| Salmeterol | 10.32 | ±0.18 | 57.1 | ±3.5 | 9.85 | ±0.21 | 32.4 | ±1.6*** |
| Isoproterenol | 7.25 | ±0.12 | 100.6 | ±1.3 | 6.15 | ±0.09** | 101.1 | ±1.2 |
| Formoterol | 9.68 | ±0.18 | 83.7 | ± 3.1 | 9.02 | ±0.21* | 78.0 | ±3.2 |
| ICI118551 | - | - | 2.7 | ±1.6 | - | - | 4.34 | ±3.6 |

Data parameters are presented as mean ± s.e.m and are from 5 different experiments per environment. For ICI118551, the effect at 10 μM antagonist is recorded as R<sub>max</sub>. Significant differences between pEC<sub>50</sub> or R<sub>max</sub> data in the two buffers are indicated by \* P<0.05, \*\* P<0.01, \*\*\* P<0.001 (unpaired Student's t-test).

**Table 4:** Agonist potencies and maximal responses derived from TMR-GαS19cha18 binding using endpoint or kinetically derived data from high sodium experiments.

| Compounds | Endpoint |  |  |  | Kinetics rate |  |  |  |
| --- | --- | --- | --- | --- | --- | --- | --- | --- |
|  | pEC<sub>50</sub> | ±s.e.m (M) | R<sub>max</sub> | ±s.e.m (%) | pEC<sub>50</sub> | ±s.e.m (M) | R<sub>max</sub> | ±s.e.m (%) |
| Salbutamol | 5.91 | ±0.22 | 26.6 | ±3.4 | 5.66 | ±0.20 | 20.18 | ±2.34 |
| Salmeterol | 9.85 | ±0.21 | 32.4 | ±1.6 | 10.02 | ±0.34 | 22.28 | ±2.51 |
| Isoproterenol | 6.15 | ±0.09 | 101.1 | ±1.2 | 5.92 | ±0.10 | 102.5 | ±0.87 |
| Formoterol | 9.02 | ±0.21 | 78.0 | ±3.2 | 8.73 | ±0.16 | 74.1 | ±1.73 |
| ICI118551 | - | - | 4.34 | ±3.6 | - | - | 1.21 | ±0.59 |

Data are presented as mean ± s.e.m and are from 4-5 different experiments.

**Figure 3:**
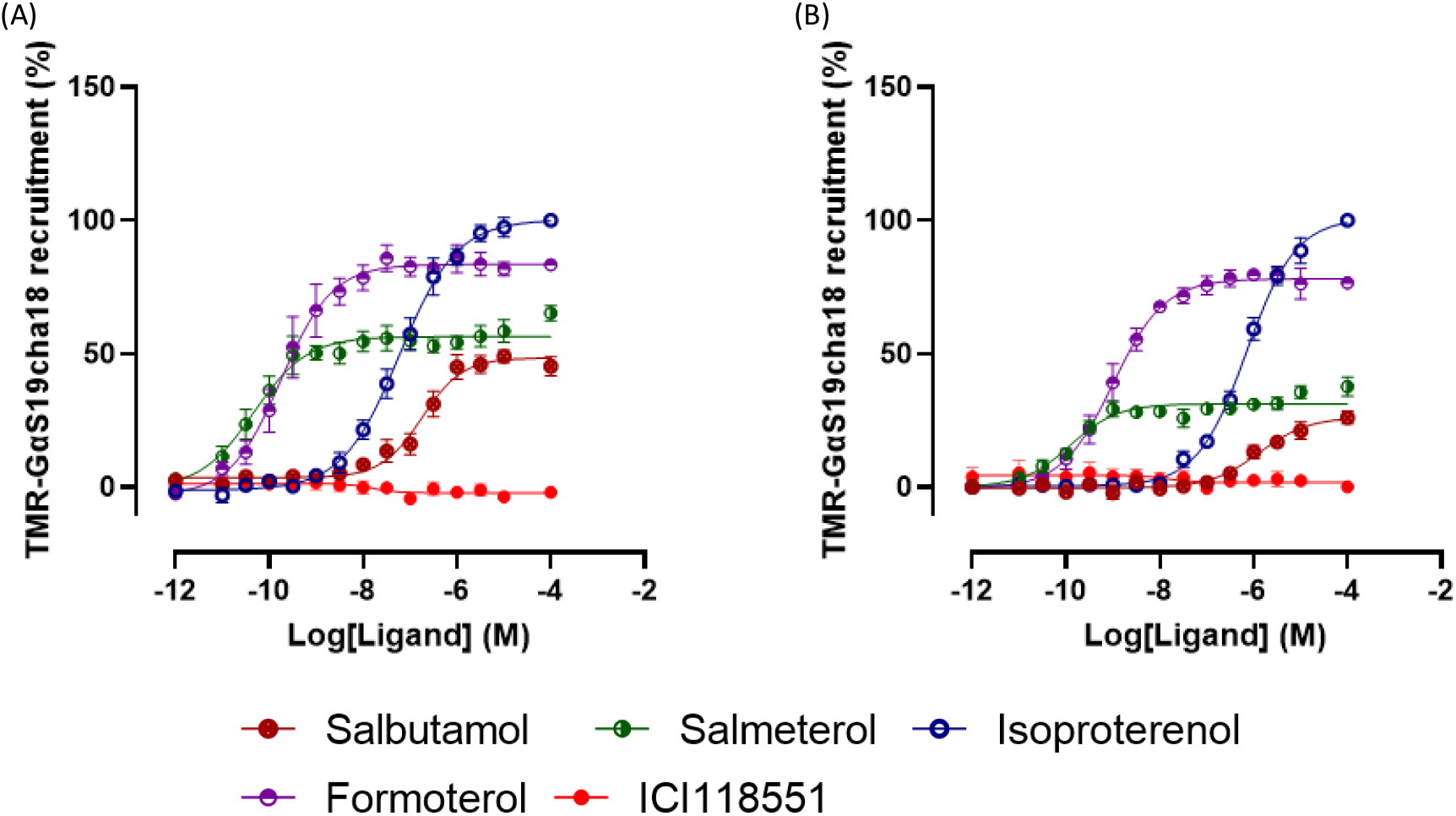
Agonist dependent recruitment of TMR-GαS19cha18 to ssβ_2_-AR-tsNluc measured by NanoBRET. Assays were performed using 500 nM TMR-GαS19cha18 with endpoint binding measured after 30 min, 37°C exposure to different β_2_-AR orthosteric ligands, to construct concentration-response relationships. A and B represent pooled data from 5 experiments, performed in low sodium and HBSS buffers respectively. In each case, agonist responses were normalized to 100 μM isoproterenol.

**Figure 4:**
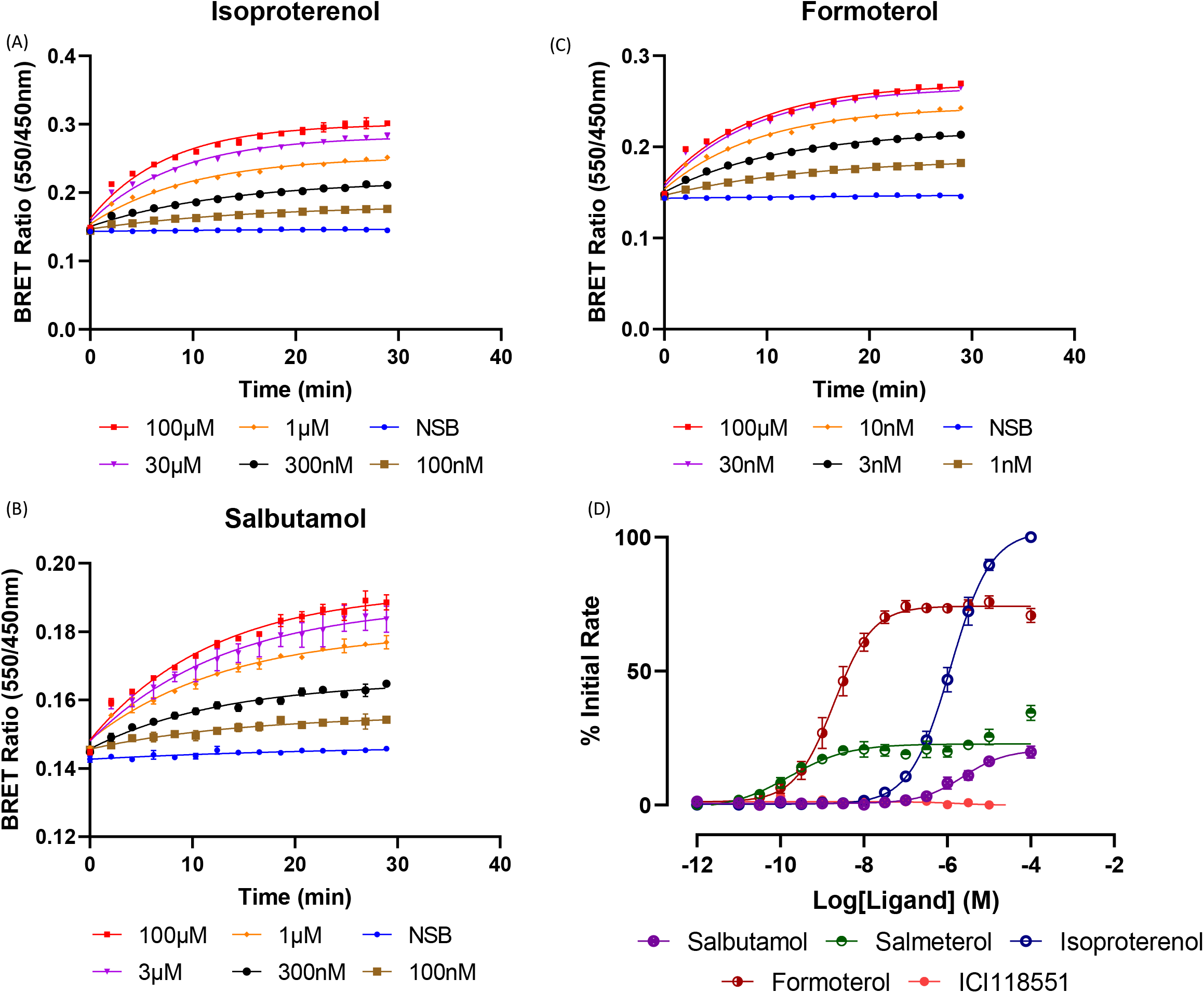
The kinetics of TMR-GαS19cha18 stimulated recruitment to the β_2_-AR. A, B and C show the concentration-dependent timecourses of TMR-GαS19cha18 recruitment measured by NanoBRET in HBSS buffer. Data are representative examples from five independent experiments. D) Initial rates of TMR-GαS19cha18 recruitment at each agonist concentration was calculated based on a rise to steady state model, and plotted to generate the pooled concentration-initial rate curves Normalized data from 4 independent experiments are shown, to the 100 μM isoproterenol response.

### TMR-GαS19cha18 is selective for Gs coupled receptors

NanoBRET binding assays employing chemokine receptor CXCR2, a Gi selective GPCR, or Gs selective prostanoid receptor EP_2_ indicated the selectivity of TMR-GαS19cha18 binding and recruitment for Gs coupled GPCRs (Figure 5). Stimulation of CXCR2-tsNluc membranes with its chemokine peptide agonist CXCL8 did not increase TMR-GαS19cha18 recruitment above basal levels. Conversely, PGE_2_ stimulation of the EP_2_-tsNluc receptor in membranes demonstrated an agonist concentration-dependent increase in TMR-GαS19cha18 NanoBRET, with levels of specific binding similar to previous β_2_-AR responses.

**Figure 5:**
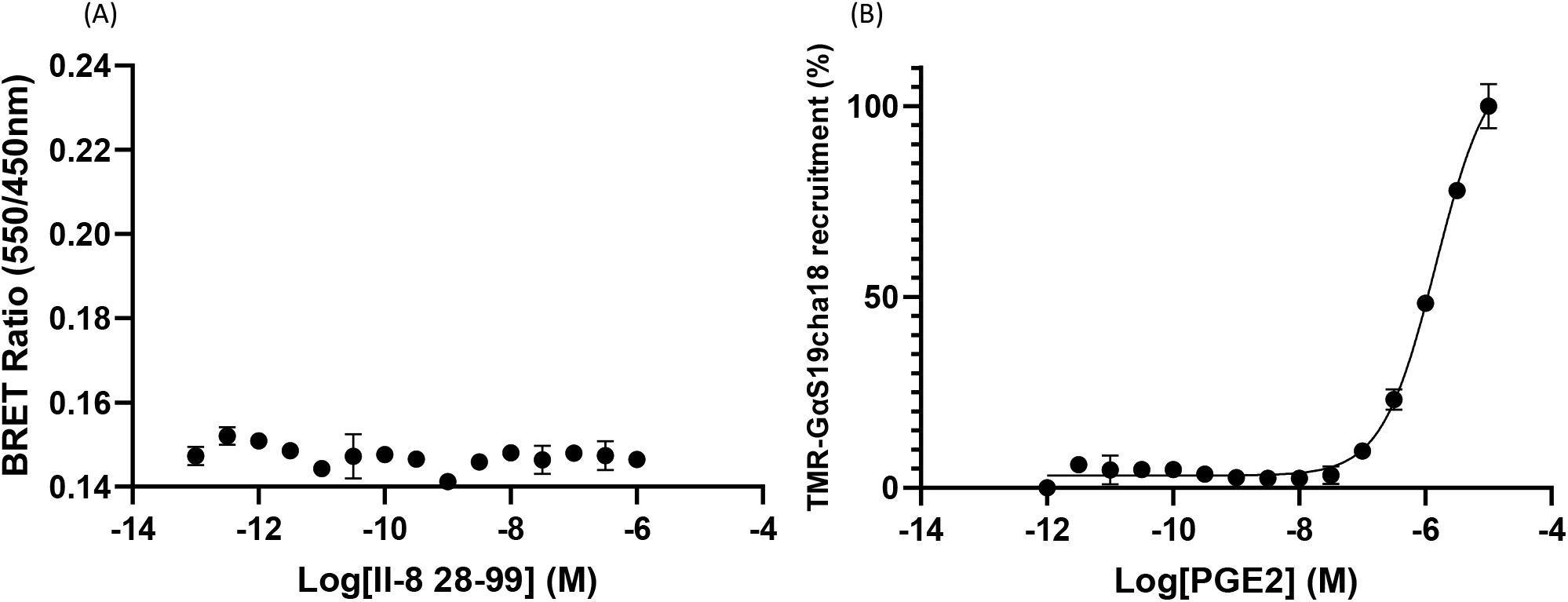
Agonist induced TMR-GαS19cha18 recruitment is also observed for the Gs coupled EP_2_ receptor but not the Gi coupled CXCR2 receptor. A) TMR-GαS19cha18 NanoBRET measurements performed in ssCXCR2-tsNLuc membranes in the absence or presence of the chemokine CXCL8 (30 min). B) Recruitment of TMR-GαS19cha18 to ssEP_2_-tsNluc measured by NanoBRET after PGE_2_ stimulation (30 min). For each receptor, data represent an individual duplicate experiment, from three performed.

### The TMR-GαS19cha18 binding assay as a detection method for ligands binding the GsGPCR G protein interaction site

To determine whether TMR-GαS19cha18could be used as a tracer in binding studies to obtain rank orders of affinity for putative IAMs, NanoBRET competition binding was performed in ssβ_2_-AR-tsNluc membranes, using the candidate unlabelled Gα C-terminal peptides GαS19cha18, GαS24, GαS11 and Gαi24 (Table 1), in the presence of isoproterenol (Figure 6). Both GαS19cha18 and GαS24 successfully competed for the G protein binding site labelled by TMR-GαS19cha18, allowing derivation of their respective affinities (GαS19cha18 > GαS24). The determined Ki for unlabelled GαS19cha18 (249 ±38nM, n=5) was equivalent to that directly measured for the TMR-GαS19cha18 probe. In contrast, GαS11 and Gαi24 did not display any detectable competition with the tracer peptide, even with a reduction in tracer concentration (Supplementary Figure 5), supporting the predicted order of selectivity of the different peptides for α5 helix binding site for Gs coupled receptors.

## Discussion

The development of selective therapeutics targeting Class A GPCRs can be limited by inherent structural homology between orthosteric binding sites for related receptor subtypes, and in this context allosteric modulation provides an attractive alternative approach to drug discovery. Within this arena, a number of successful negative intracellular allosteric modulators (IAMs) have been generated that target the receptor-G protein effector interface to inhibit signalling^8,10–14^, a mechanism that in principle should be broadly applicable to many GPCR families. The conformational selectivity of some IAMs may also be beneficial therapeutically, for example in binding the active agonist-occupied GPCR conformation preferentially to generate use dependence^1,9,34^. However, a universal route to studying and screening the G protein IAM binding site has been more challenging to identify. Here we show that Gα C-terminal peptides, which have been previously reported to act as intracellular allosteric modulators of GPCR signalling^16–18,20,21,35^, can be used as a basis to generate novel fluorescent probes for the G protein / IAM binding site, and to establish a real time, resonance energy transfer biosensor assay for binding. We demonstrate that our candidate peptide tracer acts as a novel quantitative detector of Gs receptor activation by agonists and allows development of a binding assay suitable for screening, to directly determine the affinities of competing IAM peptides and other modulators at this site.

**Figure 6:**
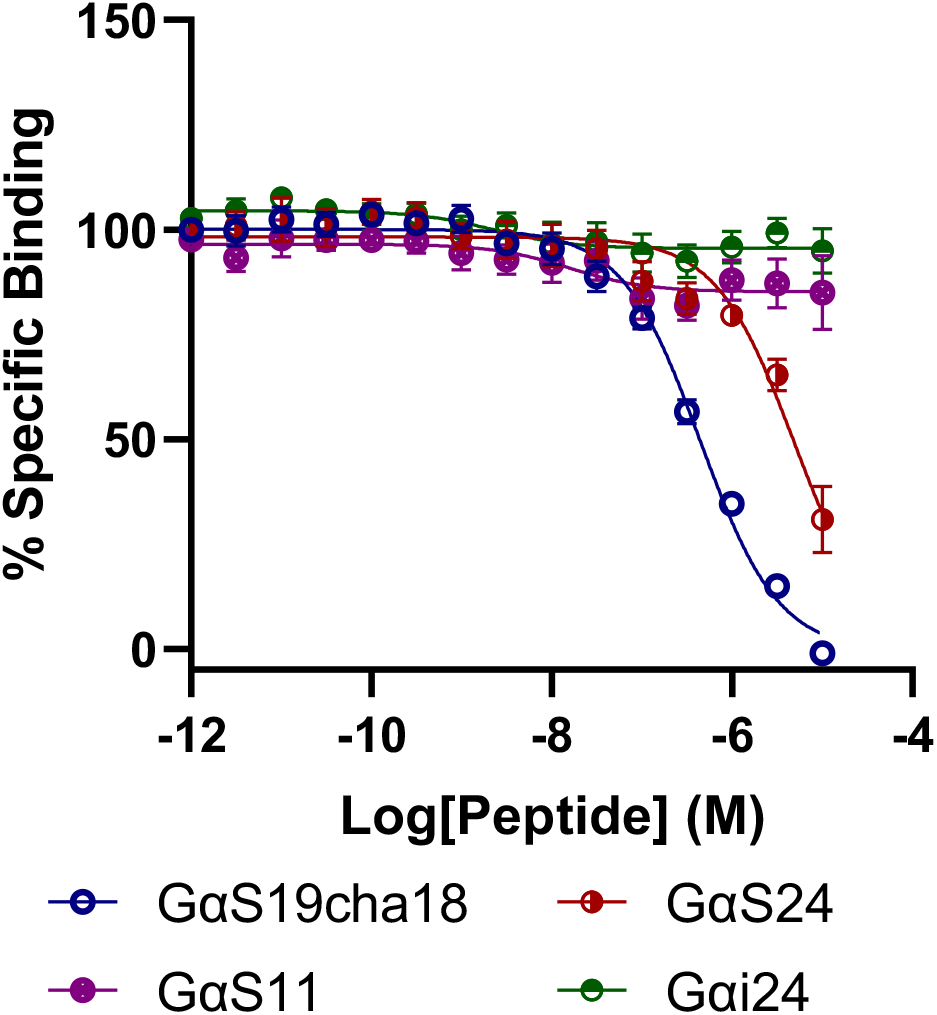
NanoBRET competition binding assays using TMR-GαS19cha18 to affinities of unlabelled Gα C terminal peptides for the ssβ_2_-AR-tsNluc receptor in membranes. Assays were performed in low sodium buffer, for 2 h at 37°C using 500 nM fluorescent tracer. Data are pooled and normalized from five independent experiments.

The length, sequence, and likely conformation of Gα C-terminal peptides have previously been shown to be essential for demonstrating binding and allosteric effects on agonist affinity^21^. We first validated these findings for key peptides using a β_2_-AR TR-FRET binding assay, using the orthosteric antagonist BODIPY-FL-propranolol^24,31^ as the fluorescent probe.

Use of a membrane assay format also allowed candidate peptides unrestricted access to the β_2_-AR intracellular surface. Although shorter C terminal Gα peptides have previously been reported to be functionally active^17,18,20^, our binding assay did not detect an ability of the 11 amino acid GαS11 to influence orthosteric ligand binding at the β_2_-AR. However, extension of the C terminal sequence to 24 amino acids in GαS24 revealed its allosteric effect, in increasing β_2_-AR agonist affinity. This may be due to the increased stability of the GαS peptide secondary structure produced by additional predicted α-helical turns, giving greater structural homology to the native G protein α5 helix. As reported by Mannes et al (2021)^21^, substitution of the penultimate leucine residue for a cyclohexylalanine residue in the 19 amino GαS19cha18 generated a peptide with the greatest allosteric effects. The affinity shift observed with GαS19cha18 was only observed with the orthosteric agonists tested, and greatest for high efficacy agonists formoterol and isoproterenol (compared to lower efficacy salbutamol) – in line with the predictions of the ternary complex model relating orthosteric agonist efficacy to the magnitude of agonist affinity changes for uncoupled and G protein coupled receptor complexes^36^. These data confirm that cha substitution in the penultimate position of the GαS C terminal sequence appears beneficial for its interactions with the β_2_-AR G protein α5 helix binding pocket sensed by the lower ends of TM3 and TM5^7^.

Using GαS19cha18 as a template, we then generated a new TMR labelled fluorescent probe to be used in conjunction with C terminal nanoluciferase fused GPCRs in NanoBRET binding assays^22,23,25,37^. The use of BRET methodology, like TR-FRET, allows for measurement of tracer recruitment with a high signal to noise ratio, and in a homogeneous assay format without separation of the bound and free ligand. Thus, it provides the opportunity to probe ligand binding kinetics as well as equilibrium measurements at the IAM G protein binding site^22,24,25^.

TMR-GαS19cha18 saturation studies using TMR-GαS19cha18 binding to β_2_-AR-Nluc membranes demonstrated clear specific binding to the receptor detected by NanoBRET, exclusively in the presence of isoproterenol agonist. These data confirm the reciprocal allosteric effects between orthosteric agonist, active receptor confirmation and the G mimetic peptides engaging the intracellular binding site^36^, illustrating the use dependent mechanism of these probes and related G protein mimetic peptides, in which binding is enhanced by receptor stimulation with orthosteric ligands. The measured K_D_ (588±19.4 nM) of TMR-GαS19cha18 was somewhat lower than that reported by Mannes et al (2021) for the unlabelled peptide in radioligand binding studies^21^, differences that may result from the level of receptor expression in the insect sf9 cell system^21^ compared to our human HEK293 cell approach. We did not observe modification of TMR-GαS19cha18 binding with inclusion of non-hydrolysable GTP analogues in the assay buffer, which might be predicted to disrupt β_2_-AR interaction with native membrane G proteins (data not shown).

The use dependent behaviour of the TMR-GαS19cha18 probe enabled its application as a signalling biosensor to discriminate orthosteric agonist efficacies at the β_2_-AR, focussing directly on receptor conformational change to the active conformation and so excluding amplification effects from downstream signalling readouts. TMR-GαS19cha18 recruitment assays defined potencies and maximal responses (full / partial) for example representative agonists of differing efficacy, in a manner comparable to previous findings^38^. The effect of high sodium concentration, as a known negative allosteric modulator of class A GPCR conformational change to an active^39,40^, demonstrated a predicted decrease in agonist potency, and enhanced the partial agonism (reduced Rmax) apparent for those ligands (salmeterol, salbutamol) with lower intrinsic efficacy. Notably, the assay enabled simple collection of kinetic TMR-GαS19cha18 recruitment data and analysis of agonist pharmacology using initial TMR-GαS19cha18 rates of association^33^, providing the opportunity to routinely monitor time dependent, as well as equilibrium agonist behaviour in one plate format.

The advantages of the TMR-GαS19cha18 biosensor in part derive from a relatively modest affinity and rapid binding kinetics, expected to follow changes in receptor conformation faithfully during activation. This may prove beneficial compared to previously reported sensors that detect the active receptor conformation with very high affinity, including miniG proteins or Nb80 nanobody recruitment, or where the sensor is tethered in close proximity to the G protein binding site through fusion to the receptor C terminus (e.g. SPASM sensors)^41,42^. Moreover, the testing of the Gs coupled EP2 receptor or Gi coupled CXCR2 within this system demonstrated TMR-GαS19cha18’s ability to bind selectively to distinct Gs coupled receptors β_2_-AR and EP2, but not to CXCR2. The ability of TMR-GαS19cha18 to bind to further Gs selective GPCRs is outside the scope of this study, however, given the shared homology with the native alpha-subunit α5 helix, these initial findings indicate the potential for such probes to recognise the G protein binding sites of a variety of Gs coupled receptors.

A key application of TMR-GαS19cha18 NanoBRET assays would be the ability to directly determine the affinities of unlabelled ligands at the G protein binding site through competition analysis, for example in the identification of new lead molecules for IAMs. Previously, such studies have only been achieved through the generation of specific radioligand IAM probes for particular receptors, such as CXCR2^43^. As proof of concept, a NanoBRET competition binding format was established for β_2_-AR-Nluc (in the presence of saturating concentrations of isoproterenol). This allowed quantitative affinity estimation for the unlabelled peptides GαS19cha18 and GαS24, and confirmed the lack of affinity of GαS11 and Gαi24 for the β_2_-AR intracellular site – dovetailing with the indirect measurements of their action on orthosteric agonist binding. One consequence of the observed probe selectivity for the agonist-occupied receptor conformation is that in future screening efforts, such binding assays are likely to reveal negative allosteric modulators with a preference for the receptor active state, which would provide them with a use-dependant mode of action^14^. This provides an additional route for therapeutic selectivity by allowing therapeutic targeting to particular regions (e.g. CNS synapses) where the target receptors are highly active, avoiding a more general inhibitory profile that might lead to undesired on target effects.

Overall, our findings demonstrate that novel GαS mimetic fluorescent probes, in combination with receptor NanoBRET technology, provide a broad strategy to monitor activation dependent changes and binding to GsPCR intracellular modulator sites. Such biosensors provide new real time readouts for orthosteric agonist activation and quantification of agonist efficacy, as well as the ability to establish NanoBRET competition binding assays to screen candidate use dependent IAMs in a medium throughput format.

## Abbreviations

(GPCR): G protein coupled receptor
(TM): Transmembrane region
(TR-FRET): Time-resolved Förster Resonance Energy Transfer
(HTRF): Homogeneous time resolved fluorescence
(BRET): Bioluminescence resonance energy transfer
(β_2_-AR): β_2_-Adrenoceptor
(Nluc): Nanoluc Nanoluciferase
(ss): SNAPtag
(TMR): Tetramethylrhodamine
(CHA): Cyclohexylalanine

## Acknowledgements

This work was funded by The British Pharmacological Society AJ Clark studentship. The Authors would like to thank Dr Desislava Nesheva for supplying CXCR2 cell membranes and Dr Nicola Dijon for providing initial EP_2_ constructs.

## Data Availability Statement

### Author contributions

JP. Farmer, ND. Holliday, SN. Mistry and CA. Laughton designed the experiments and wrote the manuscript. JP. Farmer performed and analysed the experiments.

## References

1. Rosenbaum DM, Rasmussen SGF, Kobilka BK. The structure and function of G-protein-coupled receptors. Nature 2009 459:7245. 2009;459(7245):356–363. doi:10.1038/nature08144

2. Weis WI, Kobilka BK. Structural insights into G-protein-coupled receptor activation. Current Opinion in Structural Biology. 2008;18(6):734–740. doi:10.1016/j.sbi.2008.09.010

3. Alexander S, Mathie A, Peters J. Guide to Receptors and Channels (GRAC), 5° Ed. Br J Pharmacol. Published online 2011. doi:10.1109/ISCAS.2009.5118218

4. Flock T, Hauser AS, Lund N, Gloriam DE, Balaji S, Babu MM. Selectivity determinants of GPCR-G-protein binding. Nature. 2017;545(7654):317–322. doi:10.1038/nature22070

5. Stott LA, Hall DA, Holliday ND. Unravelling intrinsic efficacy and ligand bias at G protein coupled receptors: A practical guide to assessing functional data. Biochemical Pharmacology. 2016;101:1–12. doi:10.1016/j.bcp.2015.10.011

6. Syrovatkina V, Alegre KO, Dey R, Huang XY. Regulation, Signaling, and Physiological Functions of G-Proteins. Journal of Molecular Biology. Published online 2016. doi:10.1016/j.jmb.2016.08.002

7. Rasmussen SGF, Devree BT, Zou Y, et al. Crystal structure of the β 2 adrenergic receptor-Gs protein complex. Nature. Published online 2011. doi:10.1038/nature10361

8. Liu X, Ahn S, Kahsai AW, et al. Mechanism of intracellular allosteric β2AR antagonist revealed by X-ray crystal structure. Nature. 2017;548(7668):480–484. doi:10.1038/nature23652

9. Qu C, Mao C, Xiao P, et al. Ligand recognition, unconventional activation, and G protein coupling of the prostaglandin E2 receptor EP2 subtype. Science Advances. 2021;7(14). doi:10.1126/SCIADV.ABF1268/SUPPL_FILE/ABF1268_SM.PDF

10. Andrews G, Jones C, Wreggett KA. An intracellular allosteric site for a specific class of antagonists of the CC chemokine G protein-coupled receptors CCR4 and CCR5. Molecular Pharmacology. 2008;73(3):855–867. doi:10.1124/mol.107.039321

11. Nicholls DJ, Tomkinson NP, Wiley KE, et al. Identification of a putative intracellular allosteric antagonist binding-site in the CXC chemokine receptors 1 and 2. Molecular Pharmacology. 2008;74(5):1193–1202. doi:10.1124/mol.107.044610

12. Gonsiorek W, Fan X, Hesk D, et al. Pharmacological characterization of Sch527123, a potent allosteric CXCR1/CXCR2 antagonist. Journal of Pharmacology and Experimental Therapeutics. 2007;322(2). doi:10.1124/jpet.106.118927

13. Lu H, Yang T, Xu Z, et al. Discovery of Novel 1-Cyclopentenyl-3-phenylureas as Selective, Brain Penetrant, and Orally Bioavailable CXCR2 Antagonists. Journal of Medicinal Chemistry. Published online 2018. doi:10.1021/acs.jmedchem.7b01854

14. Jiang C, Amaradhi R, Ganesh T, Dingledine R. An Agonist Dependent Allosteric Antagonist of Prostaglandin EP2 Receptors. ACS Chemical Neuroscience. 2020;11(10):1436–1446. doi:10.1021/ACSCHEMNEURO.0C00078/SUPPL_FILE/CN0C00078_SI_001.PDF

15. Scholten D, Canals M, Maussang D, et al. Pharmacological modulation of chemokine receptor function. British Journal of Pharmacology. 2012;165(6):1617–1643. doi:10.1111/j.1476-5381.2011.01551.x

16. Hamm HE, Deretic D, Arendt A, Hargrave PA, Koenig B, Hofmann KP. Site of G protein binding to rhodopsin mapped with synthetic peptides from the α subunit. Science (1979). 1988;241(4867):832–835. doi:10.1126/science.3136547

17. Rasenick MM, Watanabe M, Lazarevic MB, Hatta S, Hamm HE. Synthetic peptides as probes for G protein function. Carboxyl-terminal G alpha s peptides mimic Gs and evoke high affinity agonist binding to beta-adrenergic receptors. Journal of Biological Chemistry. 1994;269(34):21519–21525. doi:10.1016/S0021-9258(17)31835-5

18. Gilchrist A, Li A, Hamm HE. G COOH-Terminal Minigene Vectors Dissect Heterotrimeric G Protein Signaling. Science Signaling. 2002;2002(118):pl1–pl1. doi:10.1126/scisignal.1182002pl1

19. Gilchrist A, Bünemann M, Li A, Hosey MM, Hamm HE. A dominant-negative strategy for studying roles of G proteins in vivo. Journal of Biological Chemistry. 1999;274(10):6610–6616. doi:10.1074/jbc.274.10.6610

20. Gilchrist A, Vanhauwe JF, Li A, Thomas TO, Voyno-Yasenetskaya T, Hamm HE. Gα Minigenes Expressing C-terminal Peptides Serve as Specific Inhibitors of Thrombin-mediated Endothelial Activation. Journal of Biological Chemistry. 2001;276(28):25672–25679. doi:10.1074/jbc.M100914200

21. Mannes M, Martin C, Triest S, et al. Development of Generic G Protein Peptidomimetics Able to Stabilize Active State Gs Protein-Coupled Receptors for Application in Drug Discovery. Angewandte Chemie International Edition. Published online February 17, 2021:anie.202100180. doi:10.1002/anie.202100180

22. Stoddart LA, Kilpatrick LE, Hill SJ. NanoBRET Approaches to Study Ligand Binding to GPCRs and RTKs. Trends in Pharmacological Sciences. 2018;39(2):136–147. doi:10.1016/j.tips.2017.10.006

23. Soave M, Briddon SJ, Hill SJ, Stoddart LA. Fluorescent ligands: Bringing light to emerging GPCR paradigms. British Journal of Pharmacology. 2020;177(5):978–991. doi:10.1111/bph.14953

24. Sykes DA, Stoddart LA, Kilpatrick LE, Hill SJ. Binding kinetics of ligands acting at GPCRs. Molecular and Cellular Endocrinology. 2019;485:9–19. doi:10.1016/j.mce.2019.01.018

25. Stoddart LA, White CW, Nguyen K, Hill SJ, Pfleger KDG. Fluorescence-and bioluminescence-based approaches to study GPCR ligand binding. British Journal of Pharmacology. 2016;173(20):3028–3037. doi:10.1111/bph.13316

26. Sykes DA, Moore H, Stott L, et al. Extrapyramidal side effects of antipsychotics are linked to their association kinetics at dopamine D2 receptors. Nature Communications. 2017;8(1):763. doi:10.1038/s41467-017-00716-z

27. Valentin-Hansen L, Groenen M, Nygaard R, Frimurer TM, Holliday ND, Schwartz TW. The arginine of the DRY motif in transmembrane segment III functions as a balancing micro-switch in the activation of the β2-adrenergic receptor. J Biol Chem. 2012;287(38):31973–31982. doi:10.1074/JBC.M112.348565

28. Hoare BL, Kaur A, Harwood CR, et al. Measurement of non-purified GPCR thermostability using the homogenous ThermoBRET assay. bioRxiv. Published online August 17, 2020:2020.08.05.237982. doi:10.1101/2020.08.05.237982

29. Dixon AS, Schwinn MK, Hall MP, et al. NanoLuc Complementation Reporter Optimized for Accurate Measurement of Protein Interactions in Cells. ACS Chemical Biology. 2016;11(2):400–408. doi:10.1021/acschembio.5b00753

30. Baker JG. The selectivity of β-adrenoceptor antagonists at the human β1, β2 and β3 adrenoceptors. British Journal of Pharmacology. 2005;144(3):317–322. doi:10.1038/sj.bjp.0706048

31. Sykes DA, Charlton SJ. Single Step Determination of Unlabeled Compound Kinetics Using a Competition Association Binding Method Employing Time-Resolved FRET. In: Methods in Molecular Biology. ; 2018:177–194. doi:10.1007/978-1-4939-8630-9_10

32. Sykes DA, Charlton SJ. Slow receptor dissociation is not a key factor in the duration of action of inhaled long-acting β2-adrenoceptor agonists. British Journal of Pharmacology. 2012;165(8):2672. doi:10.1111/J.1476-5381.2011.01639.X

33. Hoare SRJ, Tewson PH, Quinn AM, Hughes TE, Bridge LJ. Analyzing kinetic signaling data for G-protein-coupled receptors. Scientific Reports 2020 10:1. 2020;10(1):1–23. doi:10.1038/s41598-020-67844-3

34. Hilger D, Masureel M, Kobilka BK. Structure and dynamics of GPCR signaling complexes. Nat Struct Mol Biol. 2018;25(1):4–12. doi:10.1038/S41594-017-0011-7

35. Mazzoni MR, Taddei S, Giusti L, et al. A Gα(s) carboxyl-terminal peptide prevents G(s) activation by the A(2A) adenosine receptor. Molecular Pharmacology. 2000;58(1):226–236. doi:10.1124/mol.58.1.226

36. de Lean A, Stadel JM, Lefkowitz RJ. A ternary complex model explains the agonist-specific binding properties of the adenylate cyclase-coupled beta-adrenergic receptor. Journal of Biological Chemistry. 1980;255(15):7108–7117. doi:10.1016/S0021-9258(20)79672-9

37. Stoddart LA, Johnstone EKM, Wheal AJ, et al. Application of BRET to monitor ligand binding to GPCRs. Nat Methods. 2015;12(7):661–663. doi:10.1038/NMETH.3398

38. Dijon NC, Nesheva DN, Holliday ND, Aires S, Martins M, Miguel D. Luciferase Complementation Approaches to Measure GPCR Signaling Kinetics and Bias. Methods in Molecular Biology. 2021;2268:249–274. doi:10.1007/978-1-0716-1221-7_17

39. Katritch V, Fenalti G, Abola EE, Roth BL, Cherezov V, Stevens RC. Allosteric sodium in class A GPCR signaling. Trends Biochem Sci. 2014;39(5):233–244. doi:10.1016/J.TIBS.2014.03.002

40. Agasid MT, Sørensen L, Urner LH, Yan J, Robinson C v. The Effects of Sodium Ions on Ligand Binding and Conformational States of G Protein-Coupled Receptors-Insights from Mass Spectrometry. J Am Chem Soc. 2021;143(11):4085–4089. doi:10.1021/JACS.0C11837/ASSET/IMAGES/LARGE/JA0C11837_0002.JPEG

41. Wan Q, Okashah N, Inoue A, et al. Mini G protein probes for active G protein–coupled receptors (GPCRs) in live cells. Journal of Biological Chemistry. 2018;293(19):7466–7473. doi:10.1074/jbc.RA118.001975

42. Kim K, Paulekas S, Sadler F, et al. β2-adrenoceptor ligand efficacy is tuned by a two-stage interaction with the Gαs C terminus. Proceedings of the National Academy of Sciences. 2021;118(11):e2017201118. doi:10.1073/pnas.2017201118

43. Salchow K, Bond M, Evans S, et al. A common intracellular allosteric binding site for antagonists of the CXCR2 receptor: Research paper. British Journal of Pharmacology. Published online 2010. doi:10.1111/j.1476-5381.2009.00623.x

