## Supplementary information for "Development of fluorescent peptide G Protein Coupled Receptor activation biosensors for NanoBRET characterisation of intracellular allosteric modulators"

### Supplementary information: Development of a novel G Protein-Coupled Receptor activation biosensor for high-throughput characterisation of intracellular allosteric modulators

James P Farmer, Shailesh N Mistry, Charles A Laughton, Nicholas D Holliday

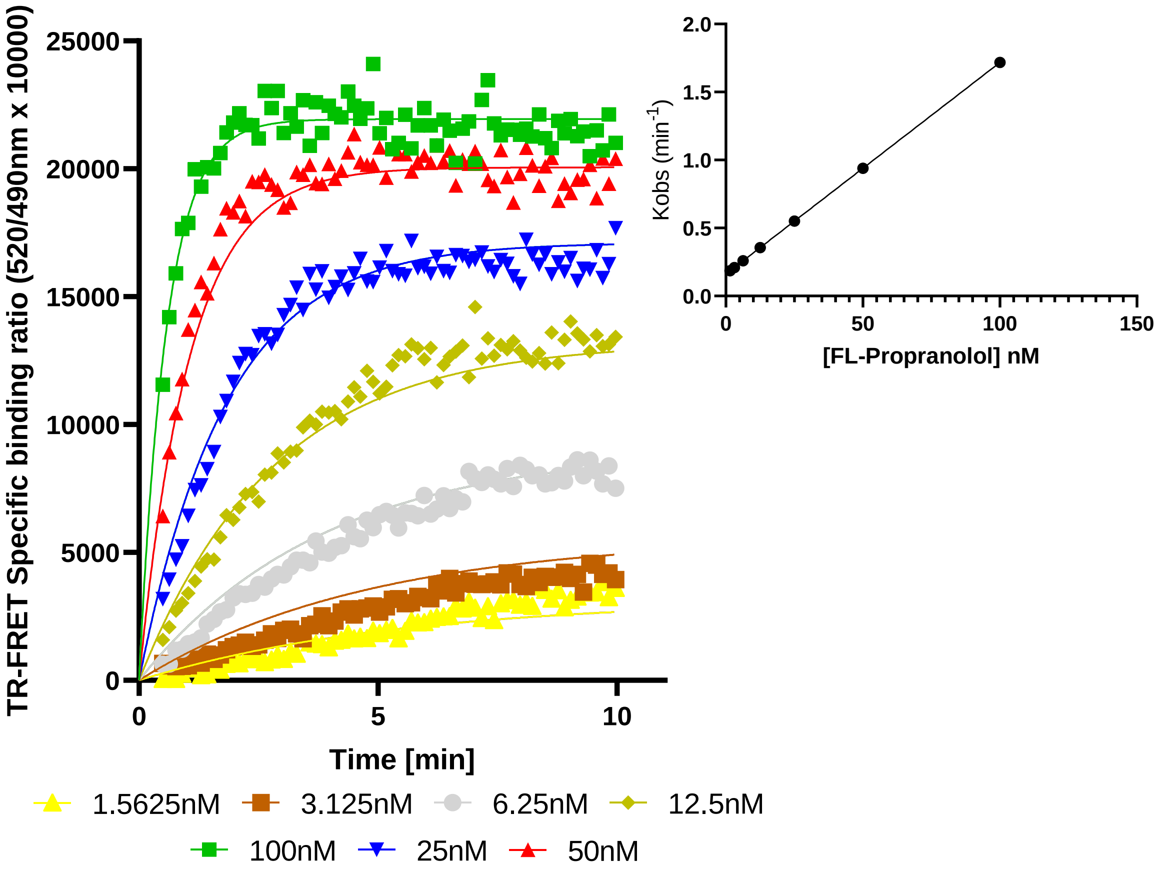


**Supplementary Figure 1: Determination of Fl-propranolol kinetic binding parameters. (left)** Fl-propranolol binding under low sodium conditions. **(right)** plot of Fl-propranolol concentration against Kobs showing binding following a simple law of mass action model with kobs increasing with concentration in a linear manner. Data in singlet from a representative of five experiments. Non-specific binding was in all cases determined through inclusion of 10µM ICI118551 and deducted from plotted data to determine specific binding.

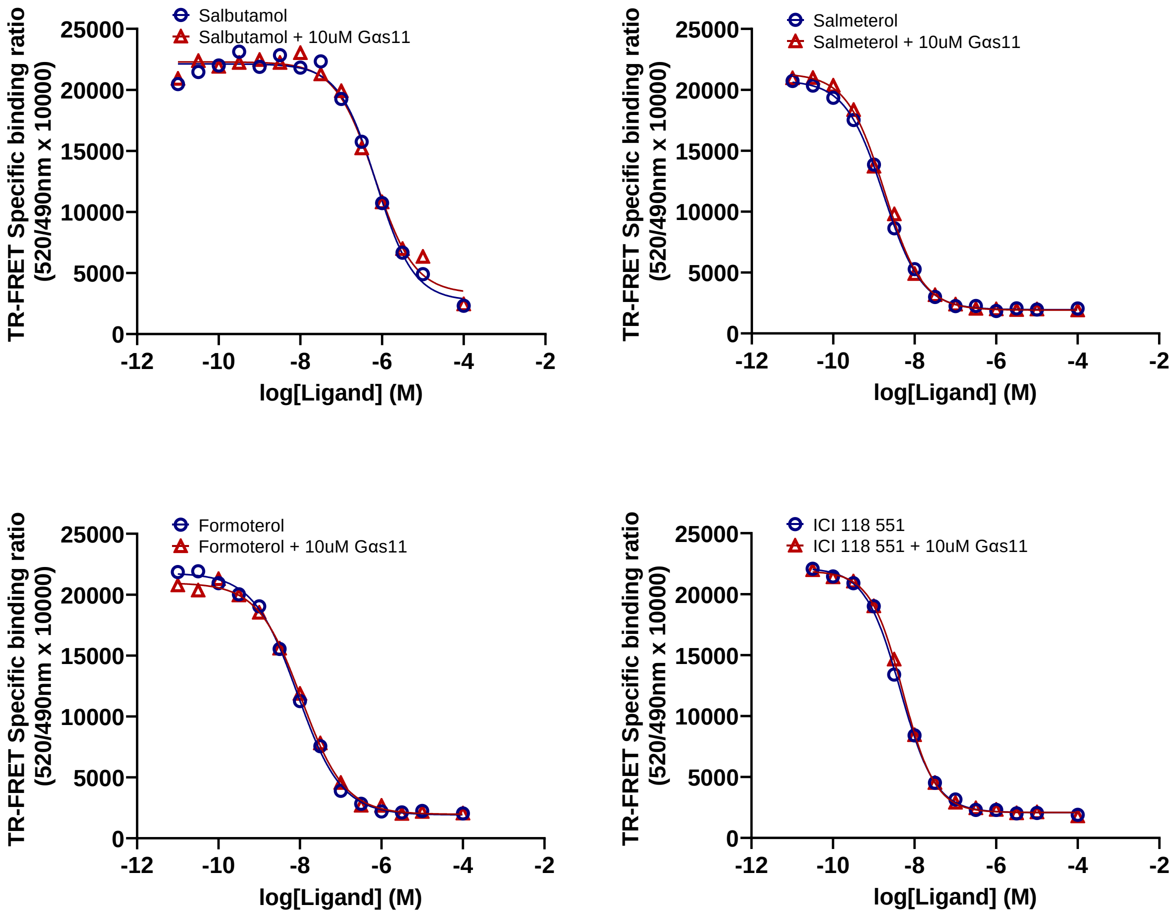


**Supplementary figure 2: Competition between 20nM Fl-propranolol and β2- adrenoceptor agonists in the presence of 10μM GαS11 in Low Sodium conditions.** Data in duplicate from a representative of five experiments. Non-specific binding was in all cases determined through inclusion of 10μM ICI118551 and deducted from plotted data to determine specific binding. All assays run at 37°C for 2 hours.

**Supplementary table 1:** Binding parameters of β2-adrenoceptor ligands in the presence of 10μM Gαs11 peptide

| **Compounds** | **pki** | **±s.e.m** |
|---|---|---|
| **Salbutamol** | **5.92** | ±0.18 |
| **Salbutamol + GαS11** | **5.78** | ±0.22 |
| **Salmeterol** | **8.74** | ±0.21 |
| **Salmeterol + GαS11** | **8.73** | ±0.19 |
| **Formoterol** | **8.05** | ±0.18 |
| **Formoterol + GαS11** | **7.95** | ±0.18 |
| **ICI 118 551** | **8.56** | ±0.12 |
| **ICI 118 551 + GαS11** | **8.53** | ±0.12 |

Data are presented as mean ± s.e.m from 5 different experiments

Test for significance using log(ki) data: * $P < 0.05$ high versus low sodium buffer, Student’s unpaired t test

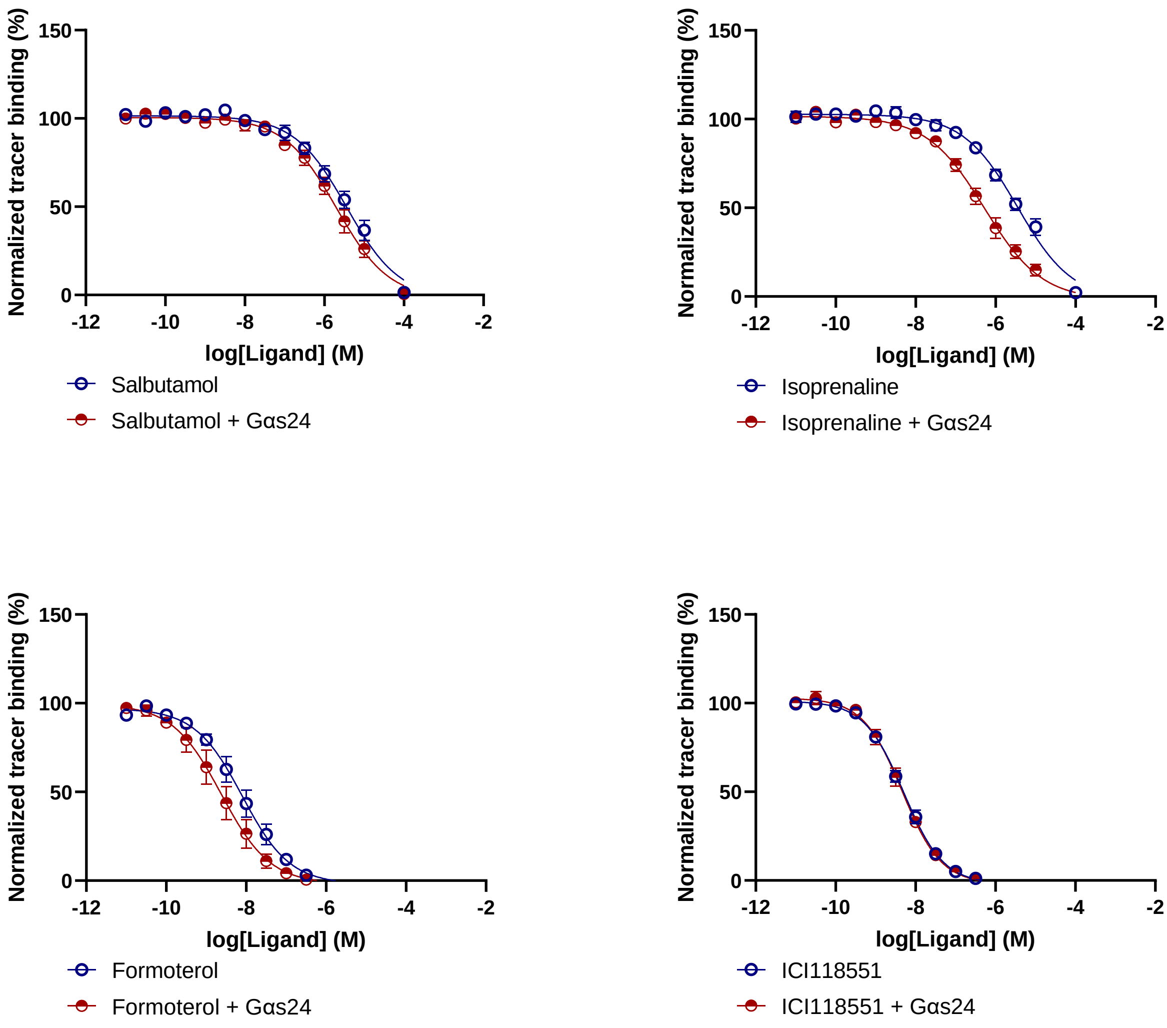


**Supplementary figure 3: Competition between 20nM Fl-propranolol and $\beta_2$- adrenoceptor agonists in the presence of 10μM GαS24 in Low Sodium conditions.** Data in duplicate from a representative of five experiments. Non-specific binding was in all cases determined through inclusion of 10μM ICI118551 and deducted from plotted data to determine specific binding. All assays run at 37°C for 2 hours.

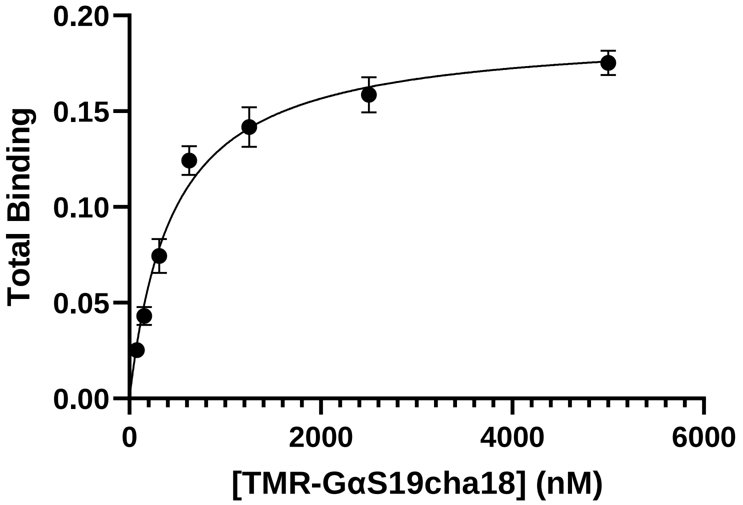


**Supplementary figure 4: Saturation binding of TMR-GαS19cha18 to the β$_2$- adrenoceptor in high sodium conditions.** Data in duplicate, taken after 60 minutes, is from a representative of three experiments. Non-specific binding was in all cases determined through inclusion of 10μM GαS19cha18 and deducted from plotted data to determine specific binding. All assays run at 37°C for 2 hours. TMR-GαS19cha18 $K_D$ = 332.8±67.58nM

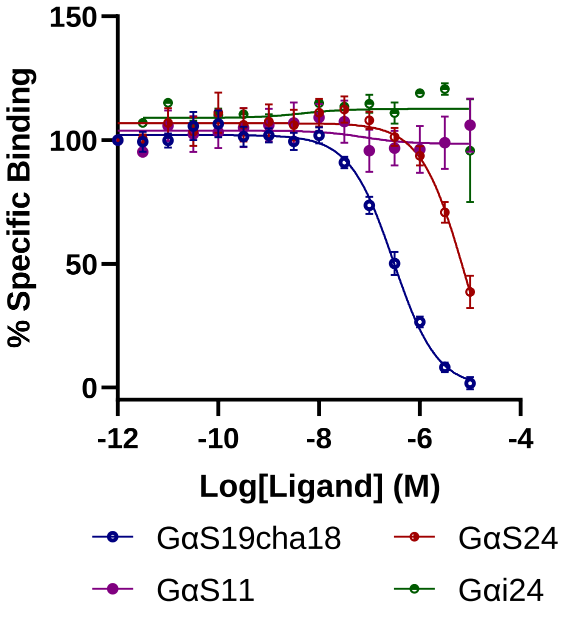


**Supplementary figure 5: Use of TMR-GαS19cha18 in competition binding assays measuring affinity of unlabelled Gα C terminal peptides for the G protein binding site.** Data is pooled, normalized data from three independent experiments. TMR-GαS19cha18 concentration reduced from 500nM to 125nM and NSB is defined by 10μM unlabelled GαS19cha18, deducted from plotted data to determine specific binding and used to determine 0%.
